# Identifying neural correlates of auditory perceptual awareness and deviance using a no-report inattentional deafness paradigm

**DOI:** 10.64898/2026.10.05.755932

**Authors:** Antonia Ellinghorst, Torge Dellert, Yunkai Zhu, Thomas Straube, Andrew R. Dykstra, Insa Schlossmacher

**Affiliations:** Institute of Medical Psychology and Systems Neuroscience, University of Münster, Münster, Germany; Otto Creutzfeldt Center for Cognitive and Behavioral Neuroscience, University of Münster, Münster, Germany; School of Communication Sciences and Disorders, University of Central Florida, Orlando, FL, USA

## Abstract

The relationship between neural processes supporting perception, deviance processing, and post-perceptual task demands during auditory processing is not well understood. We used multitone masking and a no-report inattentional deafness paradigm to dissociate neural correlates of auditory perceptual awareness, deviance processing, and task demands. Evoked responses to physically identical target-tone pairs were recorded across three phases differing only in task instructions. In Phase A, human participants remained naïve to (and thus unaware of) the target tones. In Phase B, participants were informed about (and thus aware of) the target tones. In Phase C, participants remained aware of the target tones and judged the duration of the silent gap between them, rendering them task-relevant. When participants were aware of them, target tones elicited an auditory awareness negativity and an increased positivity in the P3 window), with larger responses when target tones were additionally task-relevant. Furthermore, though largest in Phase C, deviant stimuli elicited a mismatch negativity across all task conditions, while a significant P3 to deviants was only observed in Phase C. Thus, early and late responses can be independently modulated by awareness, deviance, and task-relevance, indicating that they do not necessarily rely on shared neural mechanisms despite overlapping spatiotemporal profiles.

## Introduction

Perceiving unexpected stimuli, e.g., hearing a predator in the dark forest, is critical for survival. However, what we do know about the neural basis of conscious perception and its interaction with predictive processing is dominated by vision, leading to disproportionate reliance on this modality in theory formation (Dykstra, Cariani, et al., 2017; J. S. Snyder et al., 2015). Furthermore, there is a spatio-temporal overlap of neural correlates of awareness and mismatch processing in event-related potentials raising questions about potentially shared neural mechanisms. Understanding the interaction of conscious perception and mismatch processing in the auditory modality can thus provide novel insights into their neural basis and inform debates on competing theories.

Theories of consciousness, such as global neuronal workspace theory (GNWT; Dehaene et al., 1998, 2006; Baars, 1997, 2005), recurrent processing theory (RPT; Lamme, 2003, 2006, 2018), integrated information theory (IIT; Oizumi et al., 2014; Tononi et al., 2016), attended intermediate-level representation theory (AIR; Jackendoff, 1987; Prinz, 2012), and higher-order theories (HOT; Brown et al., 2019) propose different neural correlates of consciousness. Whereas RPT, IIT and AIR all emphasize relatively early activity in unimodal sensory association cortices, GNWT and HOT emphasize later activity in supramodal regions of frontal, parietal, and insular cortex (Förster et al., 2020; Koivisto & Revonsuo, 2010; Mashour et al., 2020), and debate persists over which of these two facets of neural activity constitutes true neural correlates of consciousness (Boly et al., 2017; Koch et al., 2016; Michel & Morales, 2019; Odegaard et al., 2017). Proponents of late activity argue that early sensory responses may be necessary but insufficient for conscious perception (Dehaene et al., 2006; Mashour et al., 2020), while proponents of early activity highlight potential confounds introduced by reporting requirements (Dembski et al., 2021; Förster et al., 2020).

Although auditory studies are comparatively limited, existing work is consistent with visual findings: both early and late correlates of awareness are observed in event-related potentials (ERPs; Dykstra, Shaw, et al., 2017; Eklund et al., 2019a, 2020; Eklund & Wiens, 2019; Filimonov et al., 2022, 2024; Sergent et al., 2021). In multitone masking paradigms, conscious perception of masker-embedded target tones is associated with activity in auditory cortex between approximately 100 and 300 ms post-stimulus (termed auditory awareness negativity, or AAN), with supramodal, P3-like activity after 300 ms (Dykstra et al., 2016; Dykstra & Gutschalk, 2015; Fernandez Pujol et al., 2023; Giani et al., 2015; Gutschalk et al., 2008). Neuroimaging and intracranial studies further implicate additional candidate regions for correlates of consciousness beyond the auditory cortex (Brancucci et al., 2011; Christison-Lagay et al., 2025; Dellert et al., 2025; Dykstra et al., 2016; Eriksson et al., 2007; Sanchez et al., 2020; Wiegand et al., 2018; Wiegand & Gutschalk, 2012).

One critical methodological issue in consciousness research concerns the confound between perceptual awareness and task demands, particularly the requirement for trial-by-trial subjective reports. Such reports can introduce task-related activity associated with maintaining information in working memory, decision making, or response preparation, potentially overestimating neural correlates of consciousness (Aru et al., 2012; Dembski et al., 2021; Förster et al., 2020; Tsuchiya et al., 2015). While several so called no-report studies exist in the visual modality (Cohen et al., 2020; Dellert et al., 2021; Hatamimajoumerd et al., 2022; Kronemer et al., 2022; Pitts et al., 2012; Schlossmacher et al., 2020; Shafto & Pitts, 2015), showing that at least some of the late responses (i.e., the P3) previously linked to perceptual awareness may instead reflect post-perceptual task demands, there are only four studies in the auditory modality (Dellert et al., 2025; Schlossmacher et al., 2021; Sergent et al., 2021; Zhu et al., 2024). Schlossmacher et al. (2021) and Zhu et al. (2024) and show that when task relevance is controlled, conscious processing of auditory speech stimuli did not elicit an increased P3 when the stimuli were not task-relevant. Furthermore, Sergent et al.(2021) showed no discernible P3 component to vowels in a passive condition, but bifurcation dynamics and late sustained negative-going activity post 300 ms that might differentiate between awareness. Interestingly, Dellert et al. (2025) showed that awareness of spoken consonants correlated with strong auditory but relatively weak fronto-parietal activations during task-irrelevant processing. These findings point to a link between P3 and postperceptual processes, although some effects in the P3 window might be observable depending on specific paradigms and analytical choices. Furthermore, past studies have been limited to speech stimuli and it remains unclear how the choice of stimuli impacts results.

In addition to their versatility in consciousness research, no-report paradigms also offer the opportunity to investigate the effects of stimulus awareness on differential processing of stimuli, such as deviants and standard stimuli in oddball designs (Dykstra & Gutschalk, 2015; Schlossmacher et al., 2020, 2022). Deviations from regular sensory input elicit characteristic brain responses, including the mismatch negativity (MMN) between ∼100 and 200 ms and a later P3 (Conroy & Polich, 2007; Näätänen et al., 2007; Polich, 2007a; E. Snyder & Hillyard, 1976; Squires et al., 1975; Stefanics et al., 2014). While the P3 is typically elicited only during conscious processing (Bekinschtein et al., 2009; Dehaene et al., 2006; Faugeras et al., 2012; Jack et al., 2017; Strauss et al., 2015), studies have observed small, MMN-like responses during nonconscious states like sleep, anesthesia, or disorders of consciousness (Koelsch et al., 2006; Strauss et al., 2015; Faugeras et al., 2012; Bekinschtein et al., 2009). However, the only study to date that used an experimental method (i.e., multitone masking) to render participants unaware produced evidence for the lack of a nonconscious MMN (Dykstra & Gutschalk, 2015). Interestingly, previous inattentional blindness studies found that the visual MMN can be elicited even if participants are unaware of the oddball stimuli (Schlossmacher et al., 2020, 2022). This might indicate that the choice of method to render stimuli unaware is important for the extent of observable differential processing (Breitmeyer, 2015). Whether similar findings can also be observed in the auditory modality using a similar methodology is yet unknown.

Here, we investigated the relationship between neural processes supporting perception, deviance processing, and post-perceptual task demands during auditory processing. We employed a three-phase multitone masking task where two target tones were reliably subliminal to naive listeners, but reliably perceived once the same listeners were informed of their presence. This design constitutes a novel auditory three-phase paradigm that, in contrast to previous work that used speech (Schlossmacher et al., 2021; Zhu et al., 2024), uses tones and inattentional deafness to hide stimuli from uninformed participants and contrast neural activity between phases in which participants were conscious and stimuli were task-irrelevant or task-relevant. To implement the oddball paradigm, we varied the stimulus onset asynchrony between the two target tones which has been shown to be able to elicit an MMN (Deouell & Bentin, 1998; Ford & Hillyard, 1981; Hari et al., 1989; Kujala et al., 2001; Näätänen et al., 1993), see Figure 1. We chose this type of deviance instead of the more conventional frequency deviant as it still allows the subjective segregation of the two target tones from the multitone masker by keeping physical features such as frequency, intensity, duration etc. constant between deviant and standard. Regarding experimental tasks, in phase A, participants completed an orthogonal decision-making task (i.e., whether more tones came from the left or right ear) while being uninformed about the target-tones (difference in gap duration). In phase B, participants were informed about the presence of the target tones, but completed the same directional decision-making task. In phase C, participants judged the duration of the silent gap between the two target tones, making them task-relevant. Thus, we were able to investigate correlates of awareness and mismatch processing while disentangling influences of awareness and task relevance.

**Figure 1.**
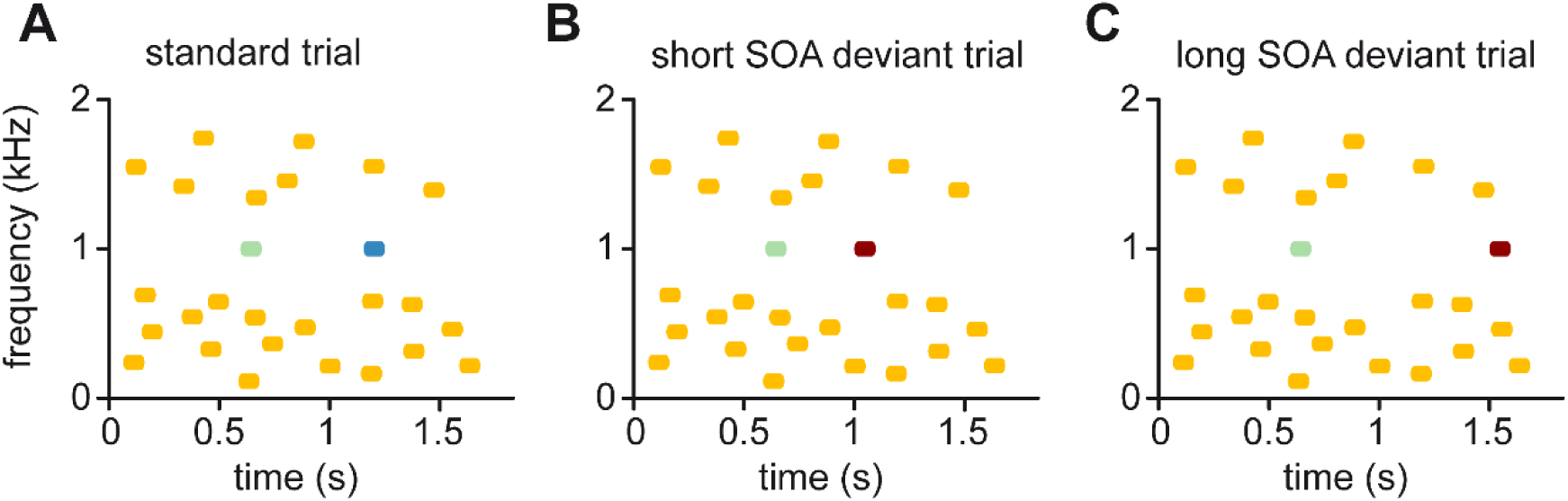
Schematic representation of different trial types. (A) Standard trial. Target tones are separated by a SOA of 600 ms. (B) Short deviant trial. Target tones are separated by a SOA of 400 ms. (C) Long SOA deviant trial. Target tones are separated by a SOA of 900 ms.

Concerning correlates of awareness, we expected that the AAN to reliably differentiate between aware and unaware target tones. Furthermore, we expected that task relevance but not awareness should be related to an increase in late positivities. Concerning mismatch processing, we expected a MMN independent of task condition, i.e. even when oddball stimuli remained unaware, while an increased P3 for deviants vs. standards should only be observed during task relevance.

## Results

Participants who reported hearing the target tones in Phase A (n = 8), indicating conscious perception from the beginning, were excluded from the EEG analysis as well as participants who reported not hearing the target tones in Phase B (n = 2) and participants whose performance measured by d’ was less than 1 in Phase C (n = 3), indicating insufficient perception of target tones. Furthermore, participants with less than 60 % of trials in EEG analysis or more than 10 interpolated electrodes were excluded (n = 2). In total, 15 participants were excluded resulting in a total of 27 participants.

### Behavioral data

The ANOVA on task performance in Phases A and B showed a significant main effect of ITD condition (*F*_8,208_ = 780.43, *p* <.001, *ɛ̂* = 0.22, BF_incl_ = 6.05×10^303^), but no significant main effect of phase (*F*_1,26_ = 0.002, *p* = 0.96, BF_excl_ = 14.13) and no interaction (*F*_8,208_ = 1.82, *p* = 0.11, *ɛ̂* = 0.62, BF_excl_ = 59.38). In Phase C, *d’* significantly differed from zero (*t*_26_ = 21.99, *p* < 0.001, BF_10_ = 1.71×10^15^). See Figure 2A and B for a visualization of task performance.

**Figure 2.**
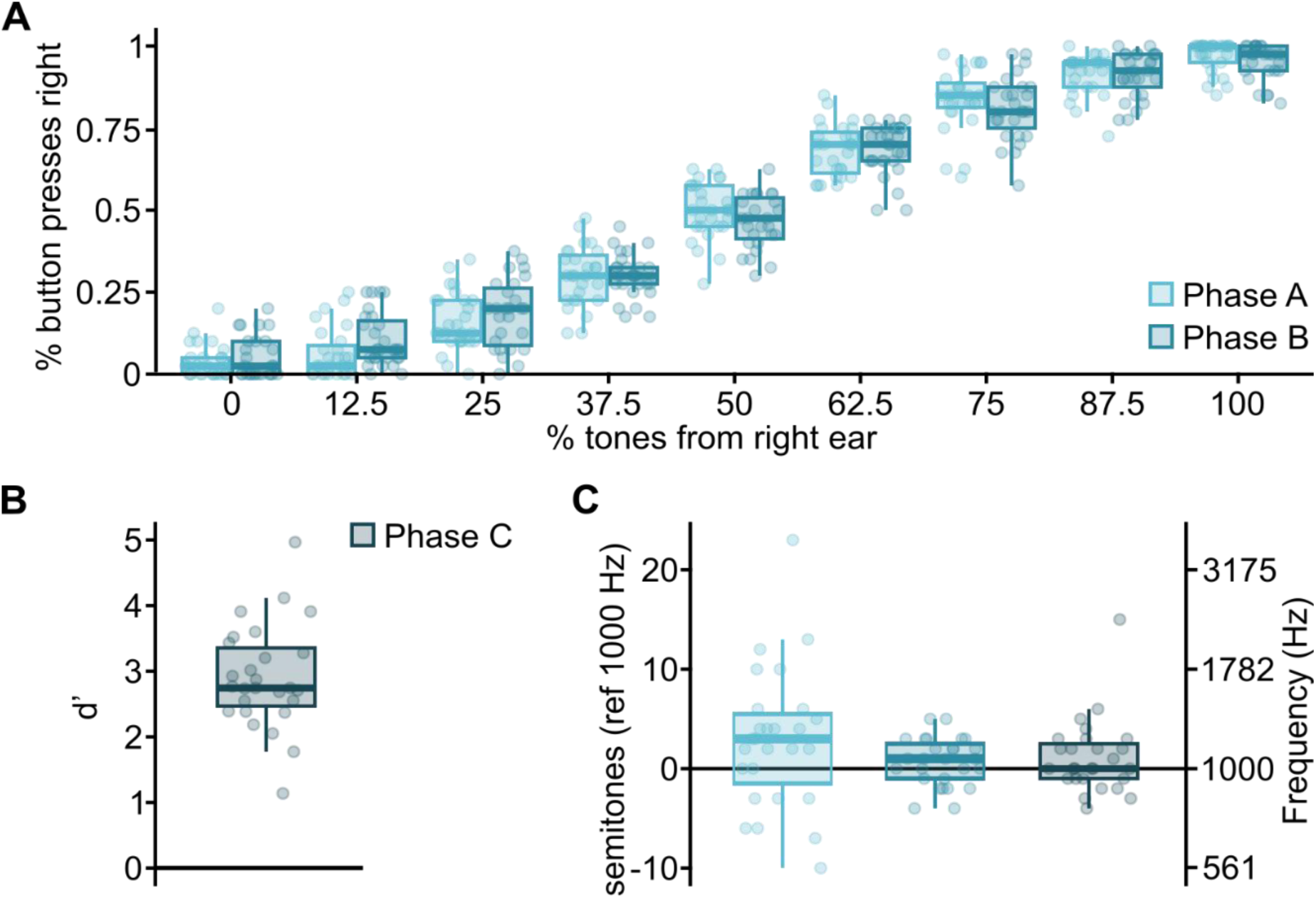
Behavioral results. (A) Performance in Phases A and B in the tone direction task. The percentage of button presses indicating more tones coming from the right side is plotted in relation to the percentage of tones presented to the right ear. (B) Performance measured as d’ in phase C in the SOA task. (C) Performance in the pitch matching task. Boxplots illustrate the variance in semitone steps from 1000 Hz. Values closer to zero indicate better performance.

In the pitch matching task, Mauchly’s test for sphericity on the number of semitone steps from 1000 Hz revealed a significant effect (*w* = 0.67, *p* = 0.007). Variance in Phase A was larger than in Phase B (*ψ̂* = 46.07, *p_holm_* < 0.001) and Phase C (*ψ̂* = 38.54, *p_holm_* = 0.03), while variance in Phases B and C did not differ (*ψ̂* = −7.53, *p_holm_* = 0.36), see Figure 2C.

### EEG data

#### Awareness and task relevance

In the AAN time window, there was one significant cluster extending from 100 to 200 ms (sum(*F*) = 10314.85, *p* < 0.001) (Figure 3A). Post hoc, Holm-Bonferroni corrected *t*-tests on cluster averages indicated a significant difference between Phases A and B (*t*_26_ = −3.32, *p* = 0.003, BF_10_ = 14.38), as well as between Phases B and C (*t*_26_ = −6.45, *p* < 0.001, BF_10_ = 22537.84).

**Figure 3.**
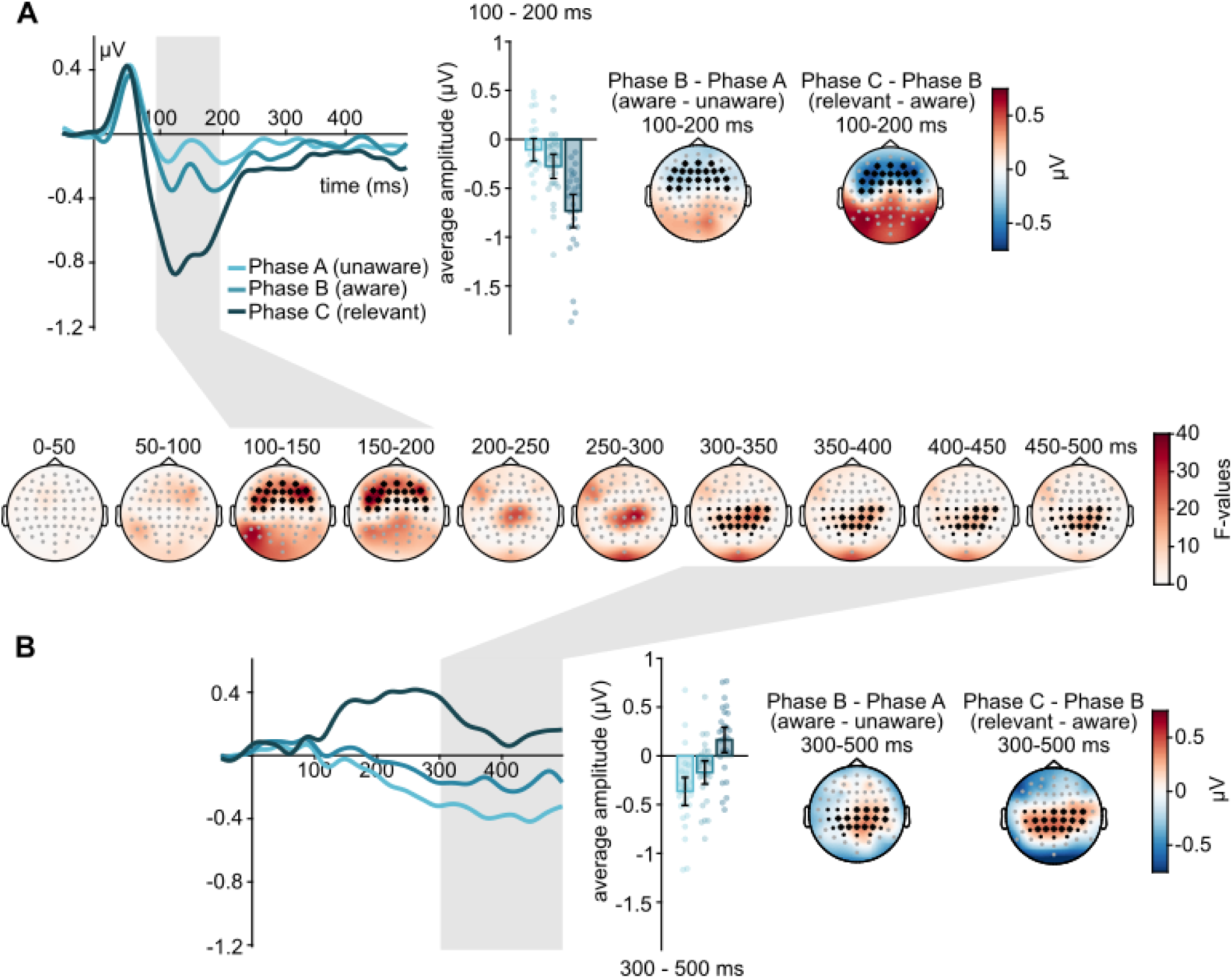
Waveforms, bar plots and topographies of effects of awareness and task relevance. (A) Early time window. (B) Late time window. Electrodes marked in black were included in the cluster-based permutation test, while those marked in gray were not; electrodes that formed part of a significant cluster are marked by asterisks. Shaded gray areas represent time windows of significant clusters. Additionally, the time course of *F*-values is displayed in the middle. Single dots represent single subject averages in the time window and electrodes of the significant cluster.

In the P3 time window, there was one significant cluster extending from 300 to 500 ms (sum(*F*) = 5495.86, *p* < 0.001) (Figure 3B). Post hoc Holm-Bonferroni corrected *t*-tests on cluster averages indicated a significant difference consisting of a relative positivity in Phase B compared to A (*t*_26_ = 2.84, *p* = 0.009, BF_10_ = 5.25), and Phase C compared to Phase B (*t*_26_ = 5.20, *p* < 0.001, BF_10_ = 1149.62).

Investigating the two target tones separately, results were similar to the averaged analysis. For both first and second target tone, in the AAN time window, significant clusters emerged and post hoc Holm-Bonferroni corrected *t*-tests indicated significant AANs as well as enhanced amplitudes due to task relevance. In the P3 time window, post-hoc Holm-Bonferroni corrected *t*-tests in significant clusters indicated a significant awareness effect for tone 2 but not tone 1 despite descriptively similar differences and a significant enhancement due to task relevance for both tones (see Supplement).

#### Mismatch

In the MMN time window, one significant cluster extended from 126 to 173 ms (cluster 1: sum(*t*) = −302.78, *p* < 0.001), and another extended from 188 to 200 ms (cluster 2: sum(*t*) = −62.26, *p* = 0.004) (Figure 4A). Post hoc Holm-Bonferroni corrected *t*-tests on cluster averages indicated a significant difference between deviants and standards in Phase A (*t*_26_ = −2.91, *p* = 0.01, BF_10_ = 6.05), Phase B (*t*_26_ = −2.23, *p* = 0.03, BF_10_ = 1.68), and Phase C (*t_26_* = −6.22, *p* < 0.001, BF_10_ = 1.29×10^4^) for cluster 1 (126 - 173 ms), while in cluster 2 (188 - 200 ms), a significant difference between deviants and standards was only observed in Phase C (*t_26_* = −7.77, *p* = 0.001, BF_10_ = 4.62×10^5^) but not in Phase A (*t_26_* = −1.85, *p* = 0.15, BF_01_ = 1.12) or Phase B (*t_26_* = −0.79, *p* = 0.44, BF_01_ = 3.69).

**Figure 4.**
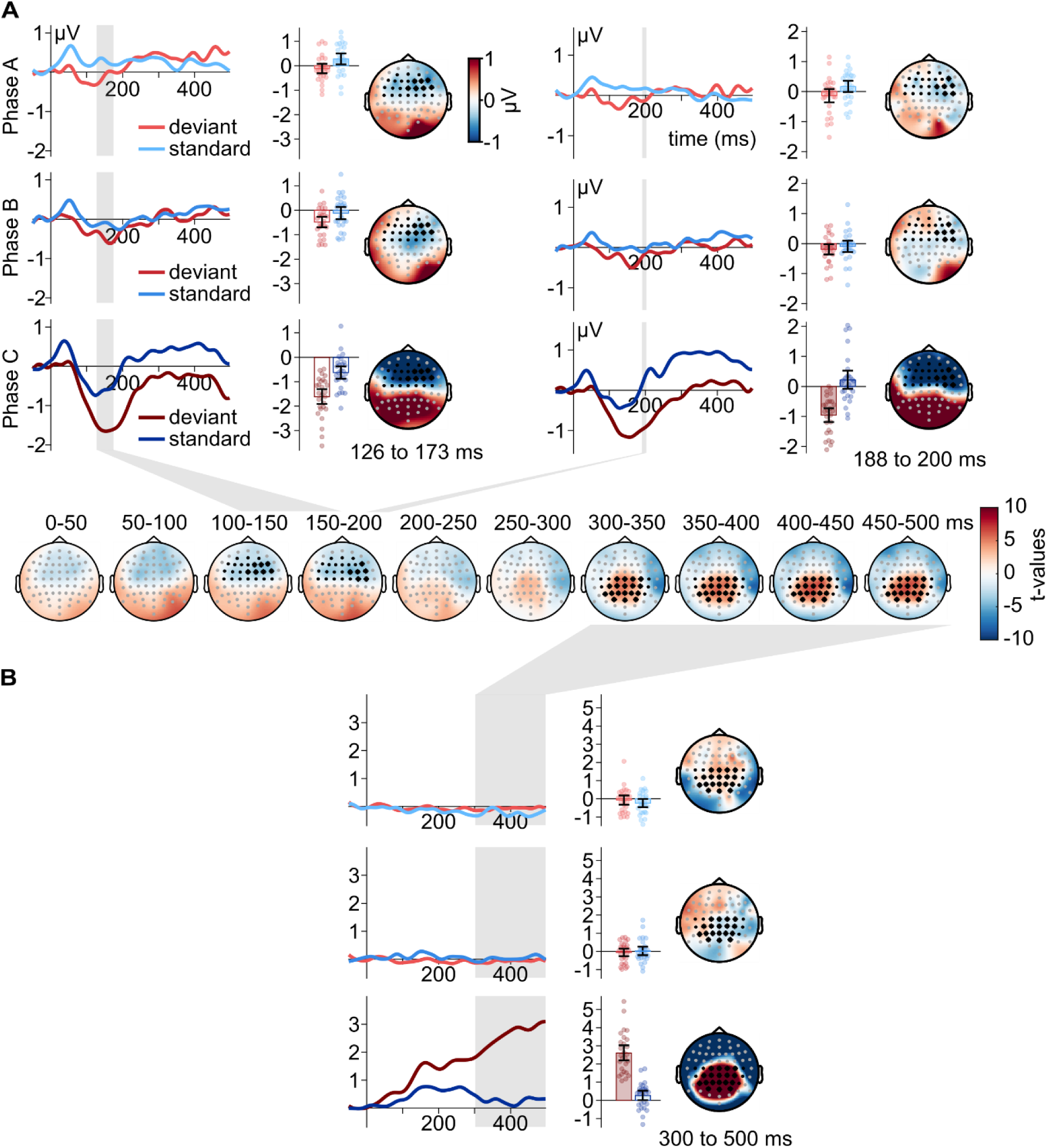
Waveforms, bar plots and topographies of effects of mismatch. (A) Early MMN time window. (B) Late P3 time window. Electrodes marked in black were included in the cluster-based permutation test, while those marked in gray were not; electrodes that formed part of a significant cluster are marked by asterisks. Shaded gray areas represent time windows of significant clusters. Additionally, the time course of *t*-values is displayed in the middle. Single dots represent single subject averages in the time window and electrodes of the significant cluster.

In the P3 time window, cluster-based permutation tests uncovered one significant cluster extending from 300 to 500 ms (sum(*t*) = 3418.61, *p* < 0.001). Post-hoc Holm-Bonferroni corrected *t*-tests on cluster averages indicated a significant difference between deviants and standards only in Phase C (*t_26_* = 10.82, *p* = 0.001, BF_10_ = 2.35×10^8^) but not in Phase A (*t_26_* = 1.10, *p* = 0.56, BF_01_ = 2.84) or Phase B (*t*_26_ = −0.48, *p* = 0.63, BF_01_ = 4.41).

## Discussion

We aimed to shed light on neural correlates of perceptual awareness and deviance (and their interaction) while controlling for task-relevance and report. Behaviorally, we manipulated participants’ awareness of otherwise salient target stimuli across phases, as evidenced by their subjective and objective reports across Phases A, B, and C (e.g., improved pitch-matching performance in Phases B and C compared to Phase A). We therefore treat target tones in Phase A as unperceived, Phase B as perceived but task-irrelevant, and Phase C as perceived and task-relevant. When the two-tone target sequence was collapsed across both tones, we found that the AAN and a late relative positivity differentiated between perceived and unperceived target tones (i.e., between Phases A and B). Furthermore, task relevance was associated with large increases in both the AAN and the relative positivity (i.e., between Phases B and C). Regarding mismatch processing, we found an MMN independent of task condition albeit smaller in unaware and aware but irrelevant conditions compared to the relevant condition. An increased P3 for deviants versus standards could only be observed for deviants in Phase C, when the two-tone target sequences (and indeed the SOA deviants) were task-relevant.

Regarding the AAN, we found increased negative amplitudes in Phase B (aware, irrelevant) compared to Phase A (unaware, irrelevant), indicating that it at least partially reflects perceptual awareness. Analysis of the two target tones separately corroborated this result with a significant difference between Phase A and B for both tones. However, amplitudes in the AAN time range were also substantially larger in Phase C (aware, relevant) compared to Phase B (aware, irrelevant), in the averaged analysis as well as the separate one. There are several possible interpretations of this effect. First, because we did not have trial-by-trial reports of perceptual awareness of the two-tone target stream, it could be simply that listeners perceived more such target streams in Phase C versus Phase B (and possibly also Phase B versus Phase A). Second, it could be that when the SOA between the target tones became task-relevant (as in Phase C), attention was more strongly directed towards the target tones (Bola & Doradzińska, 2021). Finally, it could be that such directed attention towards the target tones enhanced their perception, which would be consistent with the notion of graded (as opposed to all-or-none) perceptual awareness (Andersen et al., 2016; Doll et al., 2024).

However, in this study, a positivity in the P3 window also showed a pattern similar to the AAN. Thus, it could be argued that the same considerations apply and that the current results support the P3 as an indicator of awareness (Del Cul et al., 2007; Mashour et al., 2020), contrasting with several previous studies (Cohen et al., 2020; Dellert et al., 2021, 2022; Pitts et al., 2012, 2014; Schelonka et al., 2017; Schlossmacher et al., 2020, 2021; Sergent et al., 2021; Shafto & Pitts, 2015; Zhu et al., 2024). However, we would like to note that the current study did not reveal any P3-like effect in the absolute sense in phase B, but instead a relative negativity compared to the baseline. Only the comparison between Phase A and B revealed a relative positivity which could be tentatively linked to previous results that showed strong auditory but relatively weak fronto-parietal activations during task-irrelevant processing and highlighted the dominant role of stimulus-specific sensory areas linked to the AAN as compared to fronto-parietal ones linked to the P3 (Wiegand et al., 2018; Dellert et al., 2025). Furthermore, differences between Phase A and B did only reach significance in the analysis of tone 2 but not tone 1 in the separate analysis while descriptively showing a weak increase in both cases. Concurrently, our task-relevant condition yielded a pronounced increase in P3 amplitude compared to the irrelevant conditions with a positive amplitude in the task-relevant condition. Taken together, our findings align both with theories of consciousness emphasizing relatively early activity in sensory cortices (e.g., RPT, IIT, or AIR) *and* with theories postulating a role of fronto-parietal activity (e.g., GNWT or HOT).

Related to this, our mismatch results indicated an enhanced P3 for deviants only in the task-relevant condition, and evidence for its absence was obtained when they were task-irrelevant. This contrasts with work in patients suggesting the auditory mismatch P3 as a reliable indicator of awareness (Bekinschtein et al., 2009; Faugeras et al., 2012). However, the current findings agrees with work in the visual modality using an inattentional blindness design (Schlossmacher et al., 2020). Thus, the current results indicate that both awareness and attention are necessary preconditions for a mismatch P3. Concerning earlier potentials, however, we found an MMN-like response irrespective of awareness and relevance (though significantly larger for perceived and task-relevant stimuli). This pattern again mirrors a study using similar methodology in the visual modality (Schlossmacher et al., 2020) indicating that in both vision and audition, some aspects of mismatch processing do not require awareness (Bekinschtein et al., 2009; Koelsch et al., 2006; Strauss et al., 2015; but see Dykstra & Gutschalk, 2015).

However, there is also evidence suggesting that mismatch processing for canonical frequency deviants operates on a conscious representation of the standard stream (Dykstra & Gutschalk, 2015). Like in vision, different masking techniques and degrees may permit different levels of functional processing (Breitmeyer, 2015). In the current study, inattentional deafness was induced with a relatively weak form of multitone masking, whereas the multitone masking used in prior studies may have been stronger (e.g., Dykstra & Gutschalk, 2015; Gutschalk et al., 2008), which may have impaired nonconscious mismatch processing (Dykstra & Gutschalk, 2015). For example, most prior multitone masking studies have used both target tones and masker tones that were pure, whereas the masker tones used here were amplitude modulated. Also, finally, while we used an SOA deviant, prior work suggesting the reliance of mismatch processing on conscious representations used frequency deviants (Dykstra & Gutschalk, 2015), as in canonical auditory oddball paradigms.

In addition to an early MMN-like effect in all conditions, we obtained evidence for a later MMN effect only during task relevance. Thus, while an early automatic process indexes MMN independent from task conditions, a later MMN is tied to it. Though speculative, this pattern could be related to underlying mechanisms of MMN generation, with an early part of the MMN driven by adaptation (and occurring automatically) and a later stage that possibly indexes prediction error or short-term echoic/iconic memory traces (which may not be as automatic) (Dykstra & Gutschalk, 2015; Harenbrock et al., 2026; Schlossmacher et al., 2022, 2025; Strauss et al., 2015; Sussman et al., 2014). Future studies are needed to further investigate the underlying mechanisms of mismatch effects during inattentional deafness.

While our study has many strengths like the dissociation of awareness and task relevance, there are also some limitations that should be noted. First, while we tested our hypothesis in a classical MMN time window, descriptively the MMN has a very early onset in our study (cf. Figure 4A and B). This could be tied to the type of deviant as it has been shown that SOA deviants elicit a MMN with an earlier peak latency as compared to other deviant types (Deouell & Bentin, 1998). Thus, our results of nonconscious MMN should be tested further using other types of deviants. Furthermore, while the techniques using inattention allow to directly compare influences of awareness and task relevance, inattentional blindness and deafness paradigms have also been criticized as they might allow for an under- or overestimation of awareness depending on perspective (Lamme, 2006, 2018; Mashour et al., 2020). How these concerns apply to audition is an open question that needs to be investigated. Nevertheless, given these concerns it would be desirable to use other techniques to render stimuli unaware in order to corroborate our findings.

In summary, we found that the AAN and a late positivity scaled with increasing awareness and relevance. In contrast, the early and late mismatch responses followed a diverging pattern with no clear awareness-related differences. Thus, early and late ERPs are independently modulated by awareness and deviance, indicating that they do not rely on the same neural mechanisms despite shared spatio-temporal characteristics.

## Methods

### Participants

We recruited 42 participants (20 female) aged between 20 and 29 years (mean = 23.24; standard deviation = 1.56) for the study. All had normal hearing and reported no history of neurological or psychiatric disorders. Participants were recruited in Münster and compensated 12€ per hour. Before starting, participants were given written instructions on the experimental task and the opportunity to ask further questions. Participants provided written informed consent and the ethics committee of the German Psychological Association (Deutsche Gesellschaft für Psychologie, DGPs) approved the study. After exclusion based either on conscious perception of the target tones in Phase A (n = 8), lack of conscious perception of the target tones in Phase B (n = 2), or insufficient behavioral performance in Phase C (n = 3), a total of 27 participants remained (cf. Results for more details on exclusion criteria).

### Apparatus

The experiment was conducted within MATLAB (The Mathworks, Natick, MA, USA) using the Psychophysics Toolbox (Brainard, 1997; Kleiner et al., 2007; Pelli, 1997) on a Dell_Optiplex 5080 PC (Dell, Texas, USA). Visual stimuli were presented on a Dell P2419H monitor (Dell, Texas, USA) at 60 Hz with a resolution of 1920 × 1080 pixels. Auditory stimuli were delivered through DT 990 Pro 250 Ω (Beyerdynamic, Heilbronn, Germany) headphones using the motherboard chip (Dell Inc., 032W55, Dell, Texas, USA) for sound presentation. Participants provided their responses using a standard keyboard.

### Stimulus material

The auditory stimuli comprised two different components: (1) 24 amplitude-modulated tones that served as maskers and (2) two 1000-Hz pure tones that served as targets. All tones had a duration of 100 ms with 10-ms rise and fall times using a tapered cosine (i.e., Tukey) window. The frequency of the masker tones was chosen from logarithmically spaced frequencies between 200 and 2000 Hz, excluding those frequencies within 20% of the frequency of the target tones, leading to 15 different carrier tones (200, 227, 258, 294, 334, 379, 431, 490, 557, 632, 719, 1363, 1549, 1760, 2000 Hz) that were each amplitude-modulated at a frequency of 40 Hz and a modulation depth of 100%. Onsets of the amplitude-modulated masker tones were randomized, and every trial was divided into three 600-ms segments that each contained eight masker tones with random onset times. To lateralize the percept of the multitone masker for the perceptual decision-making task used in Phases A and B (see below), we applied an interaural time difference (ITD) of either - 604.17 µs (left-leading) or +604.17 µs (right-leading) to all masker tones. 604.17 µs was chosen because it is both near the maximum ITD that occurs naturally in humans with smaller heads (and thus generates a strongly lateralized percept) and exactly equal to 29 samples at the sampling frequency used to create the stimuli (48 kHz). Trials were uniformly distributed regarding how many had left- or right-leading ITDs ranging from 0% right-leading (i.e., 100% left-leading) to 100% right-leading in steps of 12.5% (i.e., 0, 12.5, 25, 37.5, 50, 62.5, 75, 87.5, and 100% right-leading or, correspondingly, 100, 87.5, 75, 62.5, 50, 37.5, 25, 12.5, and 0% left-leading). Which tone was presented with left- or right-leading ITD was selected at random. The two target tones had a frequency of 1000 Hz and were diotic (i.e., ITD of 0 µs) and pure (i.e., no amplitude modulation). In 80% of the trials, the two target tones were presented after 600 ms and 1200 ms (SOA = 600 ms), which will hereafter be referred to as the standard SOA. The SOA in the remaining, deviant trials was either shortened to 400 ms (10%) or lengthened to 900 ms (10%).

The average sound pressure level during the experiment amounted to 71 dB (min: 62; max: 75). Each experimental condition consisted of 360 trials, including 288 trials with standard SOA, 36 trials with short SOA, and 36 trials with long SOA. Concurrently to the auditory stream, a white fixation cross was presented on a black screen, and participants were instructed to fixate it.

### Experimental Procedure

Participants were presented with identical physical stimulation across all experimental conditions, while the specific task varied. In Phase A, listeners were instructed to report on a trial-by-trial basis whether there were more tones arriving in the left or right ear. After each trial, a prompt on the screen indicated that participants should press either the right or left arrow key on the keyboard. To familiarize participants with the task, a short practice session with only two ratios (0 and 100%), was presented prior to the first experimental run. Each practice condition was repeated three times. During this practice session, feedback was provided by coloring the frame of the chosen option green or red, indicating correct or incorrect responses, respectively. In the main experimental run, participants did not receive feedback.

Unbeknownst to participants, all trials also contained two salient target tones (cf. Stimulus material) that nevertheless remained subliminal in Phase A due to inattentional deafness caused by the multitone masker and performance of the distractor task. After Phase A, listeners were queried about their perception of this hidden two-tone sequence via a questionnaire. The first question directly queried the participants whether they perceived the two clear tones. Then, participants completed a pitch-matching task which started with the presentation of two identical tones at either 840.90 Hz or 1189.21 Hz (randomly chosen) with an SOA of 600 ms. 840.90 and 1189.21 Hz are three semitones below and above the target-tone frequency of 1000 Hz, respectively. Participants could then adjust the two-tone sequence up or down (in semitone steps) with the up and down arrow keys until the sequence best matched the 1000-Hz target-tone sequence (to the best of their ability). Lastly, they were presented with a single ‘foil’ trial that only contained the amplitude modulated masker tones and queried whether they perceived the target tones in the example.

To further investigate the perception of the participants, a structured interview was done during the instructions of the following experimental phase as we noticed during piloting that a certain percentage of the participants tended to mistake higher frequency tones (> 1363 Hz) for the targets. Thus, these participants tended to answer the awareness as well as the foil question with yes despite being unaware of the two-tone target sequence. During the instruction, target tones were first played on their own then with increased volume (+10 dB) with amplitude-modulated maskers and lastly, targets and maskers were presented at the same volume. After this, participants were asked again whether they perceived the target tones in the previous phase. Based on the outcome of the structured interview participants were either treated as unaware or aware.

The task during Phase B was the same as in Phase A: decide whether there were more left- or right-sided tones. In Phase C, the two-tone target sequence was made task-relevant by asking the participants to complete a three-alternative forced-choice regarding whether the SOA between the first and the second target tone had a standard, short or long length. The order of Phases B and C was counterbalanced to mitigate the possibility of order effects. While Phase B was not prefaced by a practice session, a short practice session was done before for Phase C in which the volume of the target was increased by 10 dB. In total, 12 trials were presented with a ratio of 4 left/right tones in the masker and the ratio of 2:1:1 of standard/short/long duration SOAs. Like the practice session of Phase A, feedback was provided in the practice session but not in the main experiment. Both Phases B and C were followed by the same questionnaire as Phase A.

### EEG recording and preprocessing

A 64-channel BioSemi active electrode system (BioSemi B.V., Amsterdam, Netherlands) was used to collect electrophysiological data, with electrodes placed using the 10-20 system. Instead of ground and reference, the BioSemi EEG system uses a CMS/DRL feedback loop with two additional electrodes (for more information see: http://www.biosemi.com/faq/cms&drl.htm). The EEG was recorded at a sampling rate of 512 Hz and electrode offsets were held below 20 µV. A build-in analog anti-aliasing low-pass filter of 104 Hz was applied prior to digitization.

Preprocessing of the EEG data was performed using FieldTrip (Oostenveld et al., 2010) in MATLAB. Offline filtering of the continuous data used Butterworth filters with half-power cut-offs. Data was band-stop filtered at 49–51 Hz (roll-off: −24 dB/octave) and at harmonic frequencies (up to 199–201 Hz) to minimize power-line noise. Additionally, a 59–61 Hz band-stop filter accounting for the monitor refresh rate (60 Hz) was applied. A 0.1 Hz high-pass filter (roll-off: - 12 dB/octave) was applied to remove slow drifts. Then, the EEG signal was segmented into epochs of 50 ms before until 500 ms after onset of individual target tones. Using independent component analysis (ICA), horizontal and vertical eye movements were removed from the data. Trials containing muscle artifacts and electrode jumps were manually excluded based on visual inspection, and bad channels were interpolated. Data was re-referenced from the CMS/DRL to a common average reference. All trials were baseline-corrected using the average of a prestimulus interval from −50 to 0 ms. For the awareness and task contrasts, we averaged all standard stimuli across the first and second target tone, resulting in three waveforms for each participant. Results from an analysis of first and second tone separately can be found in the Supplement. For mismatch contrasts, to equalize the number of averaged trials, we averaged second tone deviants as well as second tone standards from trials prior to deviant trials separately for each phase, resulting in six additional waveforms for each participant. Lastly, grand mean waveforms of the averaged data were computed.

### Statistical analysis

#### Behavioral data

To compare performance in Phases A and B, we used a 2×9 repeated-measures ANOVA with the factors Phase and ITD condition. Performance in Phase C was investigated by testing d’ against zero. Mauchly’s test for sphericity followed by the percentile bootstrap approach as recommended by Wilcox (2012) was used to investigate the variance differences in semitone steps from 1000 Hz.

#### EEG data

In a first step, we used cluster-based permutation tests (Groppe et al., 2011; Maris & Oostenveld, 2007). For awareness and task-relevance contrasts we compared the ERP responses in Phases A, B and C using an ANOVA approach (see Supplement for an analysis of each target tone separately). For mismatch processing, we computed the main effect across all conditions using a cluster-based permutation relying on a one-sided paired *t*-test. To enhance the power of the cluster-based permutations, we chose specific time intervals and electrode locations based on previous research. For the analysis of the AAN and MMN, we chose a time interval from 100 ms until 200 ms in frontocentral electrodes (Dykstra, Shaw, et al., 2017; Dykstra & Gutschalk, 2015; Eklund et al., 2019b; Gutschalk et al., 2008; Schlossmacher et al., 2021). For the P3, we selected frontocentral electrodes from 300 to 500 ms (Polich, 2007b; Schlossmacher et al., 2021). Clusters were formed by two or more neighboring data points (either in time or electrode) whenever the *F*-values/*t*-values exceeded the cluster threshold (α = 0.001). The cluster mass, sum(*F*)/sum(*t*), was calculated by adding all *F*-values/*t*-values within a cluster. The number of permutations was set to 10000, and the significance value for testing the null hypothesis was α = 0.05. Prior to the analysis, ERPs were down-sampled to 256 Hz and low-pass filtered at 25 Hz (roll-off: −24 dB/octave) to further enhance statistical power (Luck, 2014). As a second step, if a significant cluster was recovered, we extracted cluster averages and performed post-hoc *t*-tests comparing Phases A and B (and Phases B and C) to delineate effects of awareness and task relevance.

Whenever sphericity was violated, the Greenhouse-Geisser correction was applied and corrected p-values as well as *ɛ̂*-values are reported below. Post-hoc *t*-tests were corrected for multiple comparisons using the Holm-Bonferroni correction. As some of our conclusions rely on null effects, we additionally report Bayes factors (BF), with BF_01_ denoting the evidence for the null hypothesis and BF_10_ the evidence for the alternative hypothesis. For Bayesian ANOVAs we report the inclusion and exclusion Bayes Factors (Keysers et al., 2020). A high *inclusion* BF indicates that models including a given predictor are more likely than models excluding it (i.e., evidence for a significant effect of the predictor), whereas a high *exclusion* BF indicates that models excluding a given predictor are more likely than models including it (i.e., evidence for the lack of an effect of the predictor). We use the conventions from Jeffreys (1961) to interpret the results of our Bayesian analyses. All cluster-based permutations were done using FieldTrip (Oostenveld et al., 2010) in MATLAB. Other statistical tests relied on the statistics program R (R Core Team, 2023).

## Data Availability

Data and code are available on the Open Science Framework accessible via https://osf.io/ftkyx/.

## Acknowledgements

We would like to thank Else Gebhardt for her help in piloting the experiment. Funded by the Deutsche Forschungsgemeinschaft (DFG, German Research Foundation) – Project number 468787028.

## Notes

https://osf.io/ftkyx/

### Competing Interest Statement

The authors have declared no competing interest.

### Summary of Updates

-corrected a typo in results section (sum(F) instead of sum(t) for mismatch CBPs) -updated methods to refer to t-values as well as F-values in CBPs

## References

Andersen, L. M., Pedersen, M. N., Sandberg, K., & Overgaard, M. (2016). Occipital MEG Activity in the Early Time Range (<300 ms) Predicts Graded Changes in Perceptual Consciousness. Cerebral Cortex, 26(6), 2677–2688. 10.1093/cercor/bhv108

Aru, J., Axmacher, N., Lam, A. T. A. D., Fell, J., Elger, C. E., Singer, W., & Melloni, L. (2012). Local Category-Specific Gamma Band Responses in the Visual Cortex Do Not Reflect Conscious Perception. Journal of Neuroscience, 32(43), 14909–14914. 10.1523/JNEUROSCI.2051-12.2012

Baars, B. J. (1997). In the Theater of Consciousness: Global Workspace Theory, A Rigorous Scientific Theory of Consciousness. Journal of Consciousness Studies, 4, 292–309.

Baars, B. J. (2005). Global workspace theory of consciousness: Toward a cognitive neuroscience of human experience. In S. Laureys (Ed.), Progress in Brain Research (Vol. 150, pp. 45–53). Elsevier. 10.1016/S0079-6123(05)50004-9

Bekinschtein, T. A., Dehaene, S., Rohaut, B., Tadel, F., Cohen, L., & Naccache, L. (2009). Neural signature of the conscious processing of auditory regularities. Proceedings of the National Academy of Sciences, 106(5), 1672–1677. 10.1073/pnas.0809667106

Bola, M., & Doradzińska, Ł. (2021). Perceptual Awareness Negativity—Does It Reflect Awareness or Attention? Frontiers in Human Neuroscience, 15. https://www.frontiersin.org/article/10.3389/fnhum.2021.742513

Boly, M., Massimini, M., Tsuchiya, N., Postle, B. R., Koch, C., & Tononi, G. (2017). Are the Neural Correlates of Consciousness in the Front or in the Back of the Cerebral Cortex? Clinical and Neuroimaging Evidence. Journal of Neuroscience, 37(40), 9603–9613. 10.1523/JNEUROSCI.3218-16.2017

Brainard, D. H. (1997). The psychophysics toolbox. Spatial Vision, 10, 433–436. 10.1163/156856897x00357

Brancucci, A., Franciotti, R., D’Anselmo, A., Penna, S. della, & Tommasi, L. (2011). The sound of consciousness: Neural underpinnings of auditory perception. Journal of Neuroscience, 31(46), 16611–16618. 10.1523/JNEUROSCI.3949-11.2011

Breitmeyer, B. G. (2015). Psychophysical “blinding” methods reveal a functional hierarchy of unconscious visual processing. Consciousness and Cognition, 35, 234–250. 10.1016/j.concog.2015.01.012

Brown, R., Lau, H., & LeDoux, J. E. (2019). Understanding the Higher-Order Approach to Consciousness. Trends in Cognitive Sciences, 0(0). 10.1016/j.tics.2019.06.009

Christison-Lagay, K. L., Khalaf, A., Freedman, N. C., Micek, C., Kronemer, S. I., Gusso, M. M., Kim, L., Forman, S., Ding, J., Aksen, M., Abdel-Aty, A., Kwon, H., Markowitz, N., Yeagle, E., Espinal, E., Herrero, J., Bickel, S., Young, J., Mehta, A., … Blumenfeld, H. (2025). The neural activity of auditory conscious perception. NeuroImage, 308, 121041. 10.1016/j.neuroimage.2025.121041

Cohen, M. A., Ortego, K., Kyroudis, A., & Pitts, M. (2020). Distinguishing the Neural Correlates of Perceptual Awareness and Postperceptual Processing. The Journal of Neuroscience, 40(25), 4925–4935. 10.1523/JNEUROSCI.0120-20.2020

Conroy, M. A., & Polich, J. (2007). Normative Variation of P3a and P3b from a Large Sample. Journal of Psychophysiology. (world). https://econtent.hogrefe.com/doi/10.1027/0269-8803.21.1.22

Dehaene, S., Changeux, J.-P., Naccache, L., Sackur, J., & Sergent, C. (2006). Conscious, preconscious, and subliminal processing: A testable taxonomy. Trends in Cognitive Sciences, 10(5), 204–211. 10.1016/j.tics.2006.03.007

Dehaene, S., Kerszberg, M., & Changeux, J.-P. (1998). A neuronal model of a global workspace in effortful cognitive tasks. Proceedings of the National Academy of Sciences, 95(24), 14529–14534. 10.1073/pnas.95.24.14529

Del Cul, A., Baillet, S., & Dehaene, S. (2007). Brain dynamics underlying the nonlinear threshold for access to consciousness. PLOS Biology, 5(10), e260. 10.1371/journal.pbio.0050260

Dellert, T., Balster, H., Schlossmacher, I., Bruchmann, M., Moeck, R., & Straube, T. (2025). Neural correlates of consciousness in an auditory no-report fMRI study. Current Biology, 35(23), 5721–5733.e3. 10.1016/j.cub.2025.10.026

Dellert, T., Krebs, S., Bruchmann, M., Schindler, S., Peters, A., & Straube, T. (2022). Neural correlates of consciousness in an attentional blink paradigm with uncertain target relevance. NeuroImage, 264C, 119679. 10.1016/j.neuroimage.2022.119679

Dellert, T., Müller-Bardorff, M., Schlossmacher, I., Pitts, M., Hofmann, D., Bruchmann, M., & Straube, T. (2021). Dissociating the Neural Correlates of Consciousness and Task Relevance in Face Perception Using Simultaneous EEG-fMRI. Journal of Neuroscience, 41(37), 7864–7875. 10.1523/JNEUROSCI.2799-20.2021

Dembski, C., Koch, C., & Pitts, M. (2021). Perceptual awareness negativity: A physiological correlate of sensory consciousness. Trends in Cognitive Sciences, 25(8), 660–670. 10.1016/j.tics.2021.05.009

Deouell, L. Y., & Bentin, S. (1998). Variable cerebral responses to equally distinct deviance in four auditory dimensions: A mismatch negativity study. Psychophysiology, 35(6), 745–754. 10.1111/1469-8986.3560745

Doll, L., Dykstra, A. R., & Gutschalk, A. (2024). Perceptual awareness of near-threshold tones scales gradually with auditory cortex activity and pupil dilation. iScience, 27(8), 110530. 10.1016/j.isci.2024.110530

Dykstra, A. R., Cariani, P. A., & Gutschalk, A. (2017). A roadmap for the study of conscious audition and its neural basis. Phil. Trans. R. Soc. B, 372(1714), 20160103. 10.1098/rstb.2016.0103

Dykstra, A. R., & Gutschalk, A. (2015). Does the mismatch negativity operate on a consciously accessible memory trace? Science Advances, 1(10), e1500677. 10.1126/sciadv.1500677

Dykstra, A. R., Halgren, E., Gutschalk, A., Eskandar, E. N., & Cash, S. S. (2016). Neural correlates of auditory perceptual awareness and release from informational masking recorded directly from human cortex: A case study. Frontiers in Neuroscience, 10. 10.3389/fnins.2016.00472

Dykstra, A. R., Shaw, M. E., & Gutschalk, A. (2017). Electrophysiological markers of auditory perceptual awareness and release from informational masking. The Journal of the Acoustical Society of America, 141(5), 3692–3692. 10.1121/1.4988042

Eklund, R., Gerdfeldter, B., & Wiens, S. (2019a). Effects of a manual response requirement on early and late correlates of auditory awareness. Frontiers in Psychology, 10, 2083. 10.3389/fpsyg.2019.02083

Eklund, R., Gerdfeldter, B., & Wiens, S. (2019b). Effects of a Manual Response Requirement on Early and Late Correlates of Auditory Awareness. Frontiers in Psychology, 10. 10.3389/fpsyg.2019.02083

Eklund, R., Gerdfeldter, B., & Wiens, S. (2020). Is auditory awareness negativity confounded by performance? Consciousness and Cognition, 83, 102954. 10.1016/j.concog.2020.102954

Eklund, R., & Wiens, S. (2019). Auditory awareness negativity is an electrophysiological correlate of awareness in an auditory threshold task. Consciousness and Cognition, 71, 70–78. 10.1016/j.concog.2019.03.008

Eriksson, J., Larsson, A., Åhlström, K. R., & Nyberg, L. (2007). Similar frontal and distinct posterior cortical regions mediate visual and auditory perceptual awareness. Cerebral Cortex, 17(4), 760–765. 10.1093/cercor/bhk029

Faugeras, F., Rohaut, B., Weiss, N., Bekinschtein, T., Galanaud, D., Puybasset, L., Bolgert, F., Sergent, C., Cohen, L., Dehaene, S., & Naccache, L. (2012). Event related potentials elicited by violations of auditory regularities in patients with impaired consciousness. Neuropsychologia, 50(3), 403–418. 10.1016/j.neuropsychologia.2011.12.015

Fernandez Pujol, C., Blundon, E. G., & Dykstra, A. R. (2023). Laminar specificity of the auditory perceptual awareness negativity: A biophysical modeling study. PLOS Computational Biology, 19(6), e1011003. 10.1371/journal.pcbi.1011003

Filimonov, D., Railo, H., Revonsuo, A., & Koivisto, M. (2022). Modality-specific and modality-general electrophysiological correlates of visual and auditory awareness: Evidence from a bimodal ERP experiment. Neuropsychologia, 166, 108154. 10.1016/j.neuropsychologia.2022.108154

Filimonov, D., Tanskanen, S., Revonsuo, A., & Koivisto, M. (2024). Is auditory awareness graded or dichotomous: Electrophysiological correlates of consciousness at different depths of stimulus processing. Consciousness and Cognition, 123, 103720. 10.1016/j.concog.2024.103720

Ford, J. M., & Hillyard, S. A. (1981). Event-Related Potentials (ERPs) to Interruptions of a Steady Rhythm. Psychophysiology, 18(3), 322–330. 10.1111/j.1469-8986.1981.tb03043.x

Förster, J., Koivisto, M., & Revonsuo, A. (2020). ERP and MEG correlates of visual consciousness: The second decade. Consciousness and Cognition, 80, 102917. 10.1016/j.concog.2020.102917

Giani, A. S., Belardinelli, P., Ortiz, E., Kleiner, M., & Noppeney, U. (2015). Detecting tones in complex auditory scenes. NeuroImage, 122, 203–213. 10.1016/j.neuroimage.2015.07.001

Groppe, D. M., Urbach, T. P., & Kutas, M. (2011). Mass univariate analysis of event-related brain potentials/fields I: A critical tutorial review. Psychophysiology, 48(12), 1711–1725. 10.1111/j.1469-8986.2011.01273.x

Gutschalk, A., Micheyl, C., & Oxenham, A. J. (2008). Neural correlates of auditory perceptual awareness under informational masking. PLOS Biology, 6(6), e138. 10.1371/journal.pbio.0060138

Harenbrock, J., Schlossmacher, I., Todd, J., & Straube, T. (2026). Mismatch negativity: A systematic review of the spatiotemporal pattern of adaptation and prediction error in MMN. Neuroscience & Biobehavioral Reviews, 182, 106534. 10.1016/j.neubiorev.2025.106534

Hari, R., Joutsiniemi, S. L., Hämäläinen, M., & Vilkman, V. (1989). Neuromagnetic responses of human auditory cortex to interruptions in a steady rhythm. Neuroscience Letters, 99(1–2), 164–168. 10.1016/0304-3940(89)90283-8

Hatamimajoumerd, E., Murty, N. A. R., Pitts, M., & Cohen, M. A. (2022). Decoding perceptual awareness across the brain with a no-report fMRI masking paradigm. Current Biology, 32(19), 4139–4149.e4. 10.1016/j.cub.2022.07.068

Jack, B. N., Widmann, A., O’Shea, R. P., Schröger, E., & Roeber, U. (2017). Brain activity from stimuli that are not perceived: Visual mismatch negativity during binocular rivalry suppression. Psychophysiology, 54(5), 755–763. 10.1111/psyp.12831

Jackendoff, R. (1987). Consciousness and the computational mind (pp. xvi, 356). The MIT Press.

Keysers, C., Gazzola, V., & Wagenmakers, E.-J. (2020). Using Bayes factor hypothesis testing in neuroscience to establish evidence of absence. Nature Neuroscience, 23(7), 788–799. 10.1038/s41593-020-0660-4

Kleiner, M., Brainard, & Pelli. (2007). What’s new in Psychtoolbox-3?

Koch, C., Massimini, M., Boly, M., & Tononi, G. (2016). Neural correlates of consciousness: Progress and problems. Nature Reviews Neuroscience, 17(5), 307–321. 10.1038/nrn.2016.22

Koelsch, S., Heinke, W., Sammler, D., & Olthoff, D. (2006). Auditory processing during deep propofol sedation and recovery from unconsciousness. Clinical Neurophysiology, 117(8), 1746–1759. 10.1016/j.clinph.2006.05.009

Koivisto, M., & Revonsuo, A. (2010). Event-related brain potential correlates of visual awareness. Neuroscience & Biobehavioral Reviews, Special Section: Developmental Determinants of Sensitivity and Resistance to Stress: A Tribute to Seymour “Gig” Levine, 34(6), 922–934. 10.1016/j.neubiorev.2009.12.002

Kronemer, S. I., Aksen, M., Ding, J. Z., Ryu, J. H., Xin, Q., Ding, Z., Prince, J. S., Kwon, H., Khalaf, A., Forman, S., Jin, D. S., Wang, K., Chen, K., Hu, C., Agarwal, A., Saberski, E., Wafa, S. M. A., Morgan, O. P., Wu, J., … Blumenfeld, H. (2022). Human visual consciousness involves large scale cortical and subcortical networks independent of task report and eye movement activity. Nature Communications, 13(1), 7342. 10.1038/s41467-022-35117-4

Kujala, T., Kallio, J., Tervaniemi, M., & Näätänen, R. (2001). The mismatch negativity as an index of temporal processing in audition. Clinical Neurophysiology, 112(9), 1712–1719. 10.1016/S1388-2457(01)00625-3

Lamme, V. A. F. (2003). Why visual attention and awareness are different. Trends in Cognitive Sciences, 7(1), 12–18. 10.1016/S1364-6613(02)00013-X

Lamme, V. A. F. (2006). Towards a true neural stance on consciousness. Trends in Cognitive Sciences, 10(11), 494–501. 10.1016/j.tics.2006.09.001

Lamme, V. A. F. (2018). Challenges for theories of consciousness: Seeing or knowing, the missing ingredient and how to deal with panpsychism. Phil. Trans. R. Soc. B, 373(1755), 20170344. 10.1098/rstb.2017.0344

Maris, E., & Oostenveld, R. (2007). Nonparametric statistical testing of EEG- and MEG-data. Journal of Neuroscience Methods, 164(1), 177–190. 10.1016/j.jneumeth.2007.03.024

Mashour, G. A., Roelfsema, P., Changeux, J.-P., & Dehaene, S. (2020). Conscious Processing and the Global Neuronal Workspace Hypothesis. Neuron, 105(5), 776–798. 10.1016/j.neuron.2020.01.026

Michel, M., & Morales, J. (2019). Minority reports: Consciousness and the prefrontal cortex. Mind & Language, 1–21. 10.1111/mila.12264

Näätänen, R., Jiang, D., Lavikainen, J., Reinikainen, K., & Paavilainen, P. (1993). Event-related potentials reveal a memory trace for temporal features. NeuroReport, 5(3), 310.

Näätänen, R., Paavilainen, P., Rinne, T., & Alho, K. (2007). The mismatch negativity (MMN) in basic research of central auditory processing: A review. Clinical Neurophysiology, 118(12), 2544–2590. 10.1016/j.clinph.2007.04.026

Odegaard, B., Knight, R. T., & Lau, H. (2017). Should a Few Null Findings Falsify Prefrontal Theories of Conscious Perception? Journal of Neuroscience, 37(40), 9593–9602. 10.1523/JNEUROSCI.3217-16.2017

Oizumi, M., Albantakis, L., & Tononi, G. (2014). From the Phenomenology to the Mechanisms of Consciousness: Integrated Information Theory 3.0. PLOS Computational Biology, 10(5), e1003588. 10.1371/journal.pcbi.1003588

Oostenveld, R., Fries, P., Maris, E., & Schoffelen, J.-M. (2010). FieldTrip: Open source software for advanced analysis of MEG, EEG, and invasive electrophysiological data. Computational Intelligence and Neuroscience, 2011. 10.1155/2011/156869

Pelli, D. G. (1997). The VideoToolbox software for visual psychophysics: Transforming numbers into movies. Spatial Vision, 10(4), 437–442. 10.1163/156856897X00366

Pitts, M. A., Martínez, A., & Hillyard, S. A. (2012). Visual Processing of Contour Patterns under Conditions of Inattentional Blindness. Journal of Cognitive Neuroscience, 24(2), 287–303. 10.1162/jocn_a_00111

Pitts, M. A., Padwal, J., Fennelly, D., Martínez, A., & Hillyard, S. A. (2014). Gamma band activity and the P3 reflect post-perceptual processes, not visual awareness. NeuroImage, 101, 337–350. 10.1016/j.neuroimage.2014.07.024

Polich, J. (2007a). Updating P300: An integrative theory of P3a and P3b. Clinical Neurophysiology, 118(10), 2128–2148. 10.1016/j.clinph.2007.04.019

Polich, J. (2007b). Updating P300: An integrative theory of P3a and P3b. Clinical Neurophysiology, 118(10), 2128–2148. 10.1016/j.clinph.2007.04.019

Prinz, J. (2012). The Conscious Brain. OUP USA.

R Core Team. (2023). R: A language and environment for statistical computing [Software]. http://www.R-project.org/

Sanchez, G., Hartmann, T., Fuscà, M., Demarchi, G., & Weisz, N. (2020). Decoding across sensory modalities reveals common supramodal signatures of conscious perception. Proceedings of the National Academy of Sciences, 117(13), 7437–7446. 10.1073/pnas.1912584117

Schelonka, K., Graulty, C., Canseco-Gonzalez, E., & Pitts, M. A. (2017). ERP signatures of conscious and unconscious word and letter perception in an inattentional blindness paradigm. Consciousness and Cognition, 54, 56–71. 10.1016/j.concog.2017.04.009

Schlossmacher, I., Dellert, T., Bruchmann, M., & Straube, T. (2021). Dissociating neural correlates of consciousness and task relevance during auditory processing. NeuroImage, 228, 117712. 10.1016/j.neuroimage.2020.117712

Schlossmacher, I., Dellert, T., Pitts, M., Bruchmann, M., & Straube, T. (2020). Differential Effects of Awareness and Task Relevance on Early and Late ERPs in a No-Report Visual Oddball Paradigm. Journal of Neuroscience, 40(14), 2906–2913. 10.1523/JNEUROSCI.2077-19.2020

Schlossmacher, I., Lucka, F., Peters, A., Bruchmann, M., & Straube, T. (2022). Effects of awareness and task relevance on neurocomputational models of mismatch negativity generation. NeuroImage, 262, 119530. 10.1016/j.neuroimage.2022.119530

Schlossmacher, I., Protmann, I., Dilly, J., Hofmann, D., Dellert, T., Peters, A., Roth-Paysen, M.-L., Moeck, R., Bruchmann, M., & Straube, T. (2025). Underlying mechanisms of visual mismatch responses – An EEG-fMRI study. iScience, 28(12), 114039. 10.1016/j.isci.2025.114039

Sergent, C., Corazzol, M., Labouret, G., Stockart, F., Wexler, M., King, J.-R., Meyniel, F., & Pressnitzer, D. (2021). Bifurcation in brain dynamics reveals a signature of conscious processing independent of report. Nature Communications, 12(1), Article 1. 10.1038/s41467-021-21393-z

Shafto, J. P., & Pitts, M. A. (2015). Neural Signatures of Conscious Face Perception in an Inattentional Blindness Paradigm. Journal of Neuroscience, 35(31), 10940–10948. 10.1523/JNEUROSCI.0145-15.2015

Snyder, E., & Hillyard, S. A. (1976). Long-latency evoked potentials to irrelevant, deviant stimuli. Behavioral Biology, 16(3), 319–331. 10.1016/S0091-6773(76)91447-4

Snyder, J. S., Yerkes, B. D., & Pitts, M. A. (2015). Testing domain-general theories of perceptual awareness with auditory brain responses. Trends in Cognitive Sciences, 19(6), 295–297. 10.1016/j.tics.2015.04.002

Squires, N. K., Squires, K. C., & Hillyard, S. A. (1975). Two varieties of long-latency positive waves evoked by unpredictable auditory stimuli in man. Electroencephalography and Clinical Neurophysiology, 38(4), 387–401. 10.1016/0013-4694(75)90263-1

Stefanics, G., Kremláček, J., & Czigler, I. (2014). Visual mismatch negativity: A predictive coding view. Frontiers in Human Neuroscience, 8. 10.3389/fnhum.2014.00666

Strauss, M., Sitt, J. D., King, J.-R., Elbaz, M., Azizi, L., Buiatti, M., Naccache, L., Van Wassenhove, V., & Dehaene, S. (2015). Disruption of hierarchical predictive coding during sleep. Proceedings of the National Academy of Sciences, 112(11). 10.1073/pnas.1501026112

Sussman, E. S., Chen, S., Sussman-Fort, J., & Dinces, E. (2014). The Five Myths of MMN: Redefining How to Use MMN in Basic and Clinical Research. Brain Topography, 27(4), 553–564. 10.1007/s10548-013-0326-6

Tononi, G., Boly, M., Massimini, M., & Koch, C. (2016). Integrated information theory: From consciousness to its physical substrate. Nature Reviews Neuroscience, 17(7), 450–461. 10.1038/nrn.2016.44

Tsuchiya, N., Wilke, M., Frässle, S., & Lamme, V. A. F. (2015). No-Report Paradigms: Extracting the True Neural Correlates of Consciousness. Trends in Cognitive Sciences, 19(12), 757–770. 10.1016/j.tics.2015.10.002

Wiegand, K., & Gutschalk, A. (2012). Correlates of perceptual awareness in human primary auditory cortex revealed by an informational masking experiment. NeuroImage, 61(1), 62–69. 10.1016/j.neuroimage.2012.02.067

Wiegand, K., Heiland, S., Uhlig, C. H., Dykstra, A. R., & Gutschalk, A. (2018). Cortical networks for auditory detection with and without informational masking: Task effects and implications for conscious perception. NeuroImage, 167, 178–190. 10.1016/j.neuroimage.2017.11.036

Wilcox, R. (2012). Comparing Two Groups. In Introduction to Robust Estimation and Hypothesis Testing (pp. 137–213). Elsevier. 10.1016/B978-0-12-386983-8.00005-6

Zhu, Y., Li, C., Hendry, C., Glass, J., Canseco-Gonzalez, E., Pitts, M. A., & Dykstra, A. R. (2024). Isolating neural signatures of conscious speech perception with a no-report sine-wave speech paradigm. Journal of Neuroscience, 44(8). 10.1523/JNEUROSCI.0145-23.2023

