## Supplement for "Identifying neural correlates of auditory perceptual awareness and deviance using a no-report inattentional deafness paradigm"

**Analysis of awareness and task relevance for each target tone separately.**

For the first target tone, there were three significant clusters in the AAN time window, one over left (sum(*F*) = 1629.10, *p* < .001, extending from 114 to 196 ms) and two over right hemispheric electrodes (cluster 1: sum(*F*) = 224.18, *p* = .003, extending from 118 to 134 ms; cluster 2: sum(*F*) = 632.70, p < .001, extending from 142 to 173 ms), see Figure 3. Post hoc Holm-Bonferroni corrected *t*-tests on cluster averages indicated a significant difference between Phases A and B (*t*(26) = -2.60, *p* = .02, BF_10_ = 3.30), as well as between Phases B and C (*t*(26) = -4.03, *p* < .001, BF_10_ = 71.89) in the left cluster, while in the two right clusters a significant difference between Phases B and C (cluster 1: *t*(26) = -3.75, *p* = .001, BF_10_ = 37.41; cluster 2: *t*(26) = -5.17, *p* < .001, BF_10_ = 1062.24) but not between Phases A and B could be observed (cluster 1: *t*(26) = -1.81, *p* = .08, BF_01_ = 1.16; cluster 2: *t*(26) = -0.84, *p* = .41, BF_01_ = 3.55). For the second target tone, there was one significant cluster in the AAN time window (sum(*F*) = 8639.67, *p* < .001, lasting from 100 to 200 ms). Post hoc Holm-Bonferroni corrected *t*-tests on cluster averages indicated a significant difference between Phases A and B (*t*(26) = -3.21, *p* = .003, BF_10_ = 11.45), as well as between Phases B and C (*t*(26) = -5.52, *p* < .001, BF_10_ = 2485.96).

In the P3 time window, there was one significant cluster for the first target tone (sum(*F*) = 522.02, p = .002, extending from 300 to 325 ms), see Figure 4. Post hoc Holm-Bonferroni corrected *t*-tests on cluster averages indicated inconclusive evidence between Phases A and B (*t*(26) = 1.51, *p* = .14, BF_01_ = 1.80), and a significant difference between Phases B and C (*t*(26) = 3.74, *p* = .002, BF_10_ = 37.12). For the second target tone, there was one significant cluster in the P3 time window (sum(*F*) = 6562.95, p < .001, extending from 300 to 500 ms). Post hoc Holm-Bonferroni corrected *t*-tests on cluster averages indicated a significant difference between Phases A and B (*t*(26) = 2.50, *p* = .02, BF_10_ = 2.72), as well as between Phases B and C (*t*(26) = 5.11, *p* < .001, BF_10_ = 929.56).

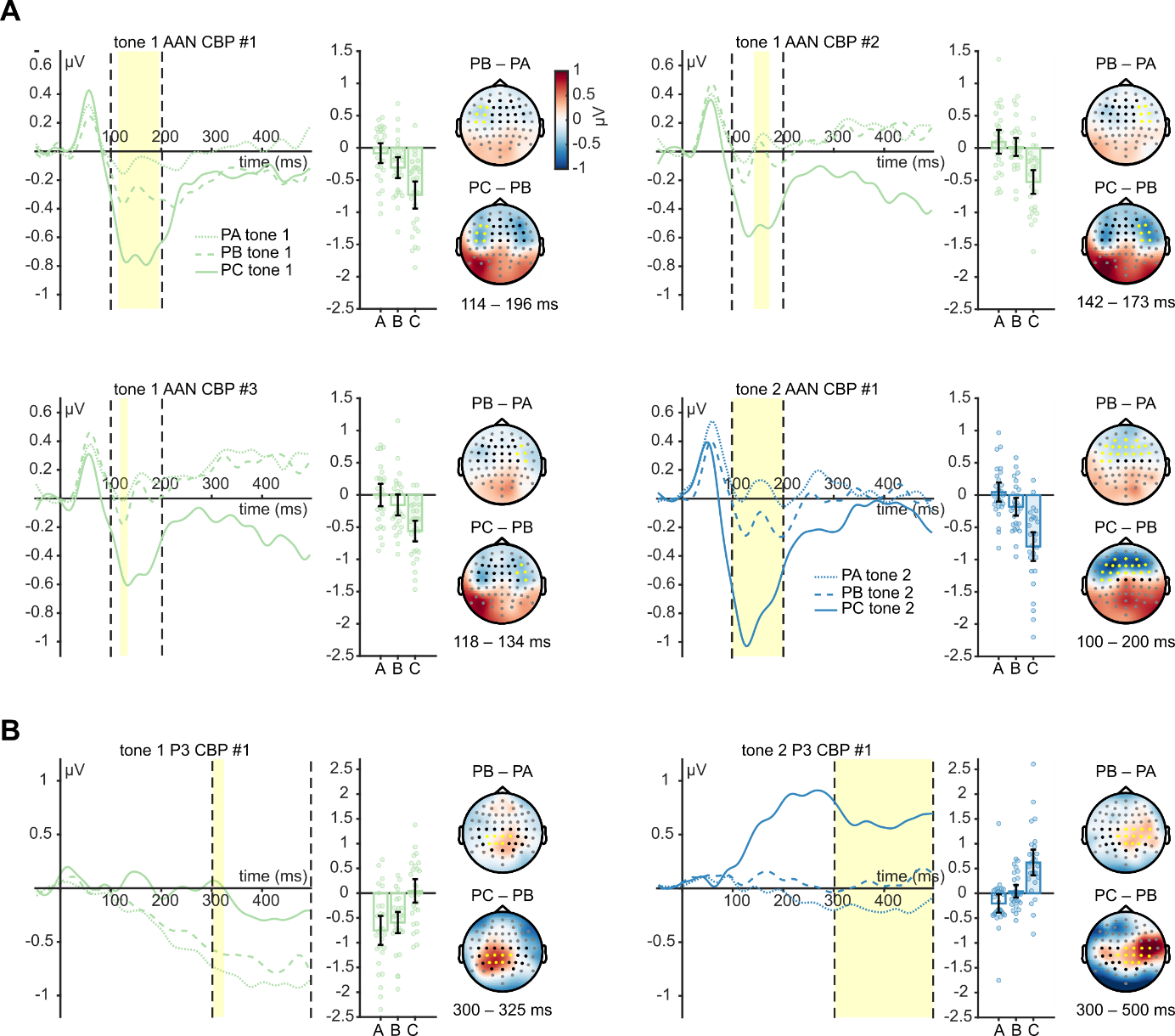

Supplementary Figure 1. Waveforms, bar plots and topographies of effects of awareness and task relevance. (A) Early time window. (B) Late time window. PA = Phase A (unaware), PB = Phase B (aware & irrelevant), PC = Phase C (aware and target-relevant). Electrodes marked in black were included in the cluster-based permutation (CBP), while those marked in gray were not; electrodes that formed part of a significant cluster are marked in yellow. Shaded yellow area represents time window of significant cluster. The first target tone is depicted in light green and the second target tone in blue. Single dots represent single subject averages in the time window and electrodes of the significant cluster.
